## Supplementary Text for "Divergence of TORC1-mediated Stress Response Leads to Novel Acquired Stress Resistance in a Pathogenic Yeast"

### Supplementary Text S1 Predicting TF binding sites in *C. glabrata* *CTA1* promoter

#### *CgCTA1* 1000bp upstream sequence

5’ -
AGCTGGTCGTTCAACTGAGAAAGTTCCAGCTTCTAAGCTATTATTTTGTCATCAAGCCTG
TGATCCCAATCTCTTTTGAGATTATCACTACTAGTTAAACACTTGTCGGAGATTAGAAGT
TTTCTCTTTCACAGGAGATATCATCTCACCAACAAAATGCTGCCTTCATCCTCTCTCTAC
CCATGTTCCTAACCTATTTGAGTTCTCTCGTTCTGTTTTGGTTCTGGGCCATGCCACAGT
ACTCTTAGTAATTGCCCTTTATCCTACTGGCTCTTGTAACCCTTCTTAACCCTTTACTAG
CTATTGCTCAAGGATAGCACAAAGGGCCATGGGATGGCTAATTCCTTAACCTTTAAAATT
TAGGGAGCCTTCCAGTTTCTCTTGTTGAATAACGGAGGTTTGTATGGAGGGAAAACGCTA
ATTTCGATGAAGGAGAGATGCCCTTCGCAAATGTGTCGAAGTTTCTCACTATTCCTTCAA
CCCTCTTTCCTCCTTTTCCTTCGGCAATAGACACTGCACTATATTTGGAGTAAAATGTCC
GTCTTTCATCGTTAAACTACCTTGAGAGAACTTAAAGAAAACTCCAACTAATTACCTTGG
AACTTGGGATAAAGAAATTAGTAACCCCTCATGCTAGTTCTCTGAAAAGTATCTGAAAAT
TTACTGTTTGCAACAATAAGTTATGGATGCAATTACAAATTGTGCTTAAGACCTAGACTA
AATTAATAACAATTATTATTTCCCTTAAGTCCTTTCTCAATAATCAATAGTTTGGATAGC
TATATAAAGGGTAAGCCTTTACCATCATCAAGCCTTGGAACACATCCTTTATCCTTTGTT
GTTCTTATTGTAAAATATATAAATACAACAACAAACCAAATCTTGAAAGCATCAATTATA
TACATAATAGATACTTTCAATCAACAAAAAATCAACATTATAACGCTAATTTTTTGTTCC
TTTTGGCTTATTAGATAACTTCCCACAATTGAAAAAAAAC

- 3’ (followed by the start codon ATG)

#### Prediction result

| **TF** | **Hit** | **% max Score** | **Position (from 3’)** | **Motif** |
| --- | --- | --- | --- | --- |
| Msn4 | CCCCT | 100 | 372 - 376 | Msn4 #518 |
| Skn7 | GGCCAT | 91.9 | 769 - 774 | Skn7 #583 |
| Skn7 | GGCCAT | 91.1 | 676 - 671 | Skn7 #583 |

#### Position frequency matrices (PFMs) used for prediction

Msn4 #518

A	0.010683761	0.002136752	0.002136752	0.002136752	0.002136752

T	0.002136752	0.002136752	0.002136752	0.002136752	0.831196581

G	0.002136752	0.002136752	0.002136752	0.002136752	0.002136752

C	0.985042735	0.993589744	0.993589744	0.993589744	0.164529915

Skn7 #583

A	0.000961538	0.000961538	0.000961538	0.000961538	0.485576923	0.208653846

T	0.000961538	0.000961538	0.000961538	0.000961538	0.000961538	0.320192308

G	0.997115385	0.997115385	0.000961538	0.1625	0.366346154	0.293269231

C	0.000961538	0.000961538	0.997115385	0.835576923	0.147115385	0.177884615

Skn7 #380 (no hit found with this motif)

A	0.054	0.084	0.032	0.054	0.202	0.035	0.101	0.022	0.108

T	0.059	0.217	0.052	0.324	0.121	0.017	0.046	0.035	0.098

G	0.84	0.279	0.042	0.309	0.363	0.876	0.428	0.03	0.2

C	0.048	0.42	0.874	0.313	0.314	0.072	0.425	0.913	0.593
